# Experience-dependent plasticity in accessory olfactory bulb interneurons following male-male social interaction

**DOI:** 10.1101/127589

**Authors:** Hillary L. Cansler, Marina A. Maksimova, Julian P. Meeks

## Abstract

Chemosensory information processing in the mouse accessory olfactory system (AOS) guides the expression of social behavior. After salient chemosensory encounters, the accessory olfactory bulb (AOB) experiences changes in the balance of excitation and inhibition at reciprocal synapses between mitral cells (MCs) and local interneurons. The mechanisms underlying these changes remain controversial. Moreover, it remains unclear whether MC-interneuron plasticity is unique to specific behaviors, such as mating, or whether it is a more general feature of the AOB circuit. Here, we describe targeted electrophysiological studies of AOB inhibitory internal granule cells (IGCs), many of which upregulate the immediate-early gene *Arc* after male-male social experience. Following the resident-intruder paradigm, *Arc*-expressing IGCs in acute AOB slices from resident males displayed stronger excitation than non-expressing neighbors when sensory inputs are stimulated. The increased excitability of *Arc*-expressing IGCs was not correlated with changes in the strength or number of excitatory synapses with MCs, but was instead associated with increased intrinsic excitability and decreased HCN channel-mediated I_H_ currents. Consistent with increased inhibition by IGCs, MCs responded to sensory input stimulation with decreased depolarization and spiking following resident-intruder encounters. These results reveal that non-mating behaviors drive AOB inhibitory plasticity, and indicate that increased MC inhibition involves intrinsic excitability changes in *Arc*-expressing interneurons.

**Significance Statement:** The accessory olfactory bulb (AOB) is a site of experience-dependent plasticity between excitatory mitral cells (MCs) and inhibitory internal granule cells (IGCs), but the physiological mechanisms and behavioral conditions driving this plasticity remain unclear. Here, we report studies of AOB neuronal plasticity following male-male social chemosensory encounters. We show that the plasticity-associated immediate-early gene *Arc* is selectively expressed in IGCs from resident males following the resident-intruder assay. After behavior, *Arc-* expressing IGCs are more strongly excited by sensory input stimulation and MC activation is suppressed. *Arc*-expressing IGCs do not show increased excitatory synaptic drive, but instead show increased intrinsic excitability. These data indicate that MC-IGC plasticity is induced after male-male social chemosensory encounters, resulting in enhanced MC suppression by *Arc*-expressing IGCs.

## Introduction

A central goal in neuroscience is to understand how sensory processing in the brain guides behavior. The mammalian accessory olfactory system (AOS) is an attractive neural pathway for studying the processes linking sensation and behavior because the AOS has a relatively linear circuit pathway into the brain and drives a variety of social behaviors (reviewed in Liberles, 2014). In the AOS, sensory information is generated in the vomeronasal organ (VNO) and processed by a single dedicated neural circuit, the accessory olfactory bulb (AOB), before being sent to limbic structures including the medial amygdala and bed nucleus of the *stria terminalis* (Scalia and Winans, 1975). Animals lacking proper AOS signaling display a range of atypical behaviors, including deficits in predator avoidance (Papes et al., 2010; Perez-Gomez et al., 2015), courtship (Pankevich et al., 2004; Kimchi et al., 2007; Ferrero et al., 2013), and territorial aggression (Stowers et al., 2002; Chamero et al., 2007; Leinders-Zufall et al., 2014). Though many of these behaviors are considered innate, behavioral and physiological evidence suggests that the AOB is a site of experience-dependent plasticity. This has been best studied in the context of pregnancy block, in which a female forms a chemosensory memory of her recent mate, but remains unexplored in the context of other AOS-mediated social behaviors (Bruce, 1959; Brennan et al., 1990).

There is still much to learn about how the AOB contributes to plasticity in social behaviors. In pregnancy block, one hypothesis proposes that inhibitory gain limits activation of AOB projection neurons, called mitral cells (MCs) to the chemosensory cues of a recent mate (reviewed in Brennan, 2009). Increased inhibition involves changes at reciprocal dendro-dendritic synapses between MCs and AOB interneurons (Brennan et al., 1990; Araneda and Firestein, 2006; Larriva-Sahd, 2008; Brennan, 2009; Smith et al., 2009). It remains unclear whether such inhibition is induced by other social chemosensory encounters, and the specific neuronal populations that contribute to these effects are not yet clear.

Immediate early genes (IEGs) are expressed in recently active neurons and can provide extensive information about the cells and networks that participate in sensory and behavioral experiences (Kawashima et al., 2013; Kim et al., 2015; Vousden et al., 2015). The IEG *Arc* is both an important plasticity-related gene and a useful marker of neuronal populations engaged by experience (reviewed in Shepherd and Bear, 2011). *Arc* is typically expressed in principal excitatory neurons (*e.g.*, cortical and hippocampal pyramidal neurons), where it mediates many forms of experience-dependent synaptic plasticity (Plath et al., 2006; Vazdarjanova et al., 2006; Jakkamsetti et al., 2013). Studies of *Arc* expression in the AOB show that it is selectively upregulated in subsets of AOB internal granule cells (IGCs), but not MCs, in male and female rodents after mating (Matsuoka et al., 2002a; Matsuoka et al., 2003). *Arc* expression by interneurons has also been noted in the main olfactory bulb in several studies (Guthrie et al., 2000; Vazdarjanova et al., 2006; Shakhawat et al., 2014). The selective expression of *Arc* by interneurons is atypical, and studying these populations is likely to provide new insights into the role of *Arc* in non-principal neuronal types.

Here, we report that *Arc* is selectively expressed in posterior AOB IGCs of male mice following the resident-intruder territorial aggression assay. *Arc* upregulation by IGCs required intact vomeronasal signaling, indicating that centrifugal inputs were not sufficient to induce *Arc* in this behavioral paradigm. Following resident-intruder behavior, *Arc*-expressing IGCs in resident males showed enhanced network excitation compared to non-expressing IGCs, while MC activity was suppressed. We investigated the IGC physiological features that underlie their enhanced activity and found no evidence for an increase in excitatory synapse strength or number. Instead, we found that *Arc*-expressing IGCs display a robust increase in intrinsic excitability compared to other IGCs. Our results show that AOB inhibitory plasticity occurs after non-mating behaviors, and reveal cellular mechanisms underlying MC inhibition after chemosensory social encounters.

## Materials and Methods

### Mice

All animal procedures were in compliance with the UT Southwestern Institutional Care and Use Committee. Sexually-naïve adult male mice aged 6-12 weeks were housed on a customized 12/12 light cycle with the lights on from noon until midnight. Food and water were provided *ad libitum*. *Arc*-*d4EGFP*-BAC (a kind gift from Pavel Osten via Kimberly Huber; Grinevich et al., 2009) and *Arc*^*tm1St*^ (“Arc-/-“ or “Arc-d2EGFP”; Jackson Labs Stock # 007662; RRID:IMSR_JAX:007662; Wang et al., 2006) mice were generous gifts from Kimberly Huber. *Trpc2*^*tm1Dlc*^ (“Trpc2-/-“ Jackson Labs Stock # 021208; RRID:IMSR_JAX:021208; Stowers et al., 2002) and *Gt(ROSA)26Sor*^*tm9(CAG-tdTomato)Hze*^ (“Ai9” Jackson Labs Stock # 007905; RRID:IMSR_JAX:007905; Madisen et al., 2010) mice were obtained from Jackson Laboratory. A total of 119 mice were used in this study.

### Behavior

Resident **male** mice were individually housed on corn cob bedding, without cage changes, for one week prior to the experiment. All behavior occurred during the dark phase (Zeitgeber time 20-24 h) in a dimly lit room to facilitate video recording. After a 10-minute habituation period, a BALB/cJ male intruder mouse was introduced to the resident cage for 10-minute encounter. To test the response to soiled bedding alone, a small petri dish was filled with bedding from a cage of 4 BALB/cJ males that had gone without cage changes for one week. This petri dish was introduced to the resident’s cage for 10 minutes instead of an intruder animal.

### Live slice preparation

Animals were anesthetized with isofluorane and decapitated 3 hours after the resident-intruder paradigm was completed unless otherwise specified. Brains were dissected and 400 µm parasagittal sections of the AOB were prepared using a Leica VT1200 vibrating microtome in ice-cold, oxygenated artificial cerebrospinal fluid (ACSF). ACSF contained (125 mM NaCl, 2.5 mM KCl, 2 mM CaCl_2_, 1 mM MgCl_2_, 25 mM NaHCO_3_, 1.25 mM NaH_2_PO_4_, 25 mM glucose, 3 mM myo-inositol, 2 mM Na-pyruvate, 0.4 mM Na-ascorbate) with an additional 9 mM MgCl_2_ in the slicing buffer. After slicing, the slices were kept in a recovery chamber at room temperature (22 °C) containing ACSF with 0.5 mM kynurenic acid to prevent potential glutamate excitotoxicity during the recovery/holding period. Just prior to recordings, slices were transferred to a slice chamber (Warner Instruments) mounted on a fluorescence-and differential interference contrast imaging-equipped upright microscope (FN1 Model, Nikon). Oxygenated ACSF was superfused via a peristaltic pump (Gilson) at a rate of 1-2 mL/min throughout. Slice temperature was maintained at 32-33 °C via inline and chamber heaters (Warner Instruments).

### 2-photon imaging and image analysis

Image stacks up to 200 µm deep were acquired using an excitation wavelength of 890 nm and a 40x (1.0 NA) water-immersion objective (Olympus). Images were denoised using a 3D median filter and deconvolved using a model point spread function in ImageJ (RRID:SCR_003070). Fluorescent cells were counted using a 3D object counting add-on (Bolte and Cordelieres, 2006). Cell counts were normalized to the volume of the cell layer of interest. *I*_*Arc*_*sum* was computed as follows:

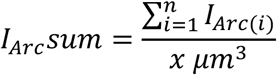

where *I*_*Arc*(*i*)_ is the normalized brightness of each cell 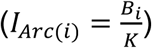 is the mean pixel intensity within cellular region of interest *i, K* is the mean pixel intensity within the ICL but outside cellular regions of interest, and *x* is the total volume within the imaged portion of the ICL. This metric combines relative brightness of all identified cells and is normalized by the imaged volume to facilitate comparisons across experimental preparations.

### Immunohistochemistry

Following behavior, animals were briefly anesthetized with inhaled isofluorane, then injected with a ketamine/xylazine cocktail (120 mg/kg ketamine/16 mg/kg xylazine dose) and transcardially perfused with 0.01 M PBS followed by 4% paraformaldehyde in PBS. Brains were post-fixed in 4% paraformaldehyde in PBS overnight. Brains were then cryoprotected overnight in PBS containing 25% sucrose, embedded in OCT compound (TissueTek), and flash frozen. 30 µm sections were prepared using a Leica CM3050 S cryostat and processed free-floating. Sections were rinsed 4x in 0.01 M PBS, incubated in 0.1% Triton X in PBS for 2 hours, rinsed 3x, incubated in 10% goat serum in PBS for 2 hours, and incubated in primary antibody in primary block (0.1% Triton X, 10% goat serum in PBS) overnight at 4 °C. Sections were then rinsed 3x in PBS and incubated in secondary antibody in secondary block (0.1% Triton X, 5% goat serum) for 2 hours. Sections were rinsed 3x, incubated in 500 nM DAPI in PBS, and rinsed 3x again. Sections were then mounted on slides in Fluoromount-G mounting medium (SouthernBiotech). Anti-Arc primary antibody specificity (Synaptic Systems #156003 rabbit polyclonal, RRID:AB_887694) was verified using *Arc*^-/-^ mice and was used at 1:1000. Anti-GFP primary (Abcam #ab13970 chicken polyclonal, RRID:AB_300798) was used at 1:500. Goat anti-rabbit AF633 (Molecular Probes Cat# A21070, RRID:AB_2535731) and goat anti-chicken AF488 (Molecular Probes Cat# A11039, RRID:AB_142924) were both used at 1:2000 dilution.

Biocytin amplification of filled neurons was achieved using streptavidin-Alexa Fluor 568 (ThermoFisher #S11226, RRID:AB_2315774). Slices were washed in PBS 3x for 30 minutes each. Slices were then incubated in a blocking solution containing 0.5% TritonX and 10% goat serum in PBS for 2 hours, followed by the same blocking solution containing streptavidin-AF568 at 0.01mg/mL for 4 hours. Slices were then rinsed 3x for 30 min each, and mounted on slides in Fluoromount-G mounting medium.

For the experiments in Figs. 2 and 5, immunostained sections were imaged with a 40x (1.3 NA) oil immersion objective on an LSM 510 inverted confocal microscope (Zeiss). For the experiments in Fig. 1, immunostained sections were imaged with a Zeiss Axioscan.Z1 using a 20x (0.8 NA) air objective. Cells were counted manually using ImageJ by a scorer blinded to the experimental condition. Anterior-posterior index was calculated using custom MATLAB software. Spines were counted and morphological analysis performed using the Simple Neuron Tracer plugin for ImageJ (Longair et al., 2011) by a scorer blinded to experimental conditions. Spines were counted on the 100 µm of primary dendrite in the external cellular layer (ECL) starting at the edge of the lateral olfactory tract.

**Figure 1:**
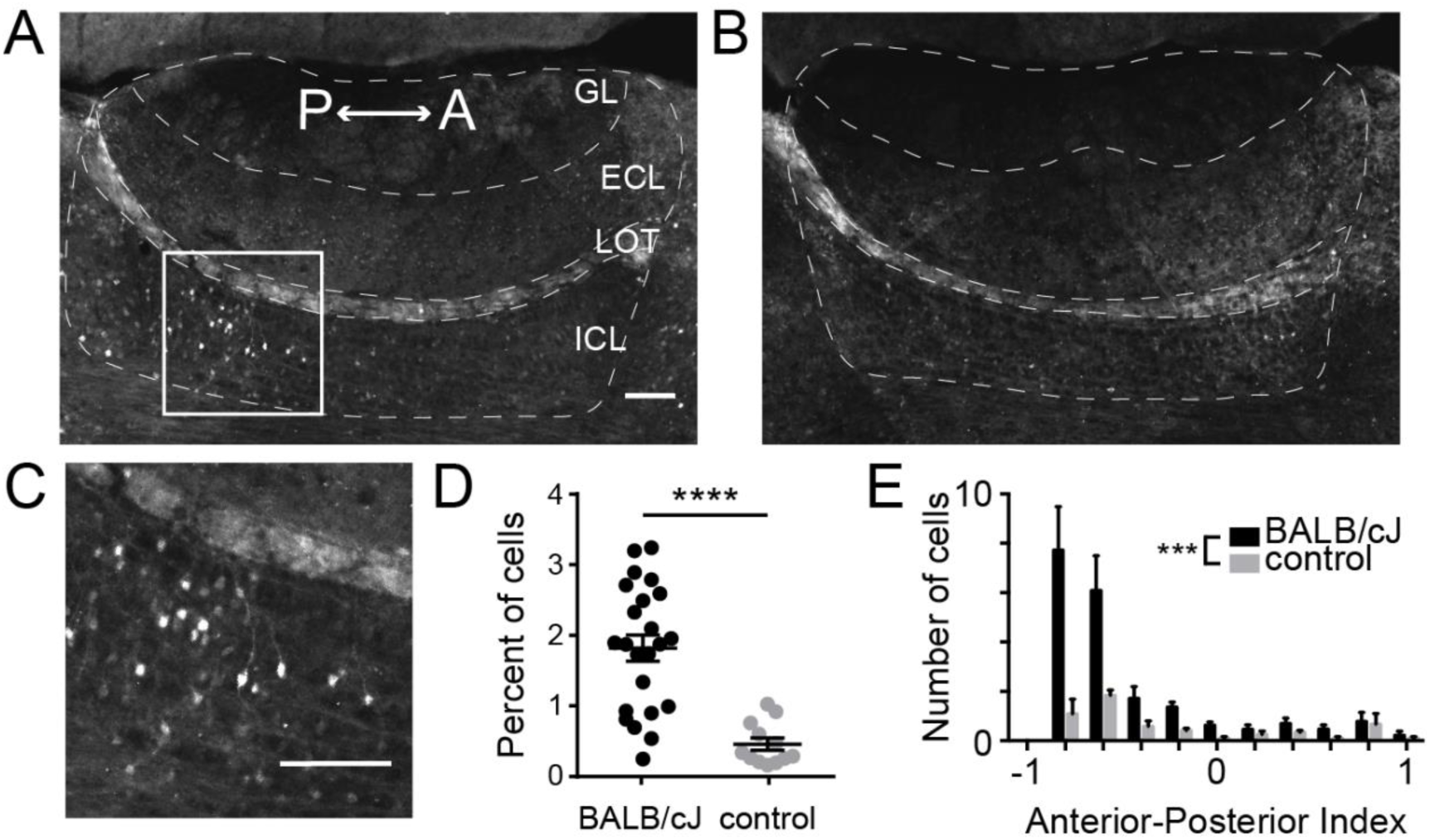
Exposing resident males to novel male intruders results in upregulation Arc protein expression in AOB IGCs. ***A,*** Parasagittal section through the AOB of a C57BL/6J male resident 90 minutes after being exposed to a BALB/cJ male intruder. Immunofluorescence indicates Arc protein expression. Scale bar: 100 µm. n=6 animals, 23 sections. GL, glomerular layer; ECL, external cellular layer; LOT, lateral olfactory tract; ICL, internal cellular layer. ***B,*** AOB section from a control resident that was not exposed to an intruder. n=3 animals, 12 sections. ***C,*** Enhanced view of IGCs in the boxed in area in panel A. ***D,*** Percentage of all cells expressing somatic Arc protein. Wilcoxon-Mann-Whitney test, **** p<0.0001. Scale bar: 100 µm. ***E,*** Normalized anterior-posterior position of Arc-expressing cells in the ICL, Wilcoxon-Mann-Whitney test, *** p<0.001.

**Figure 2:**
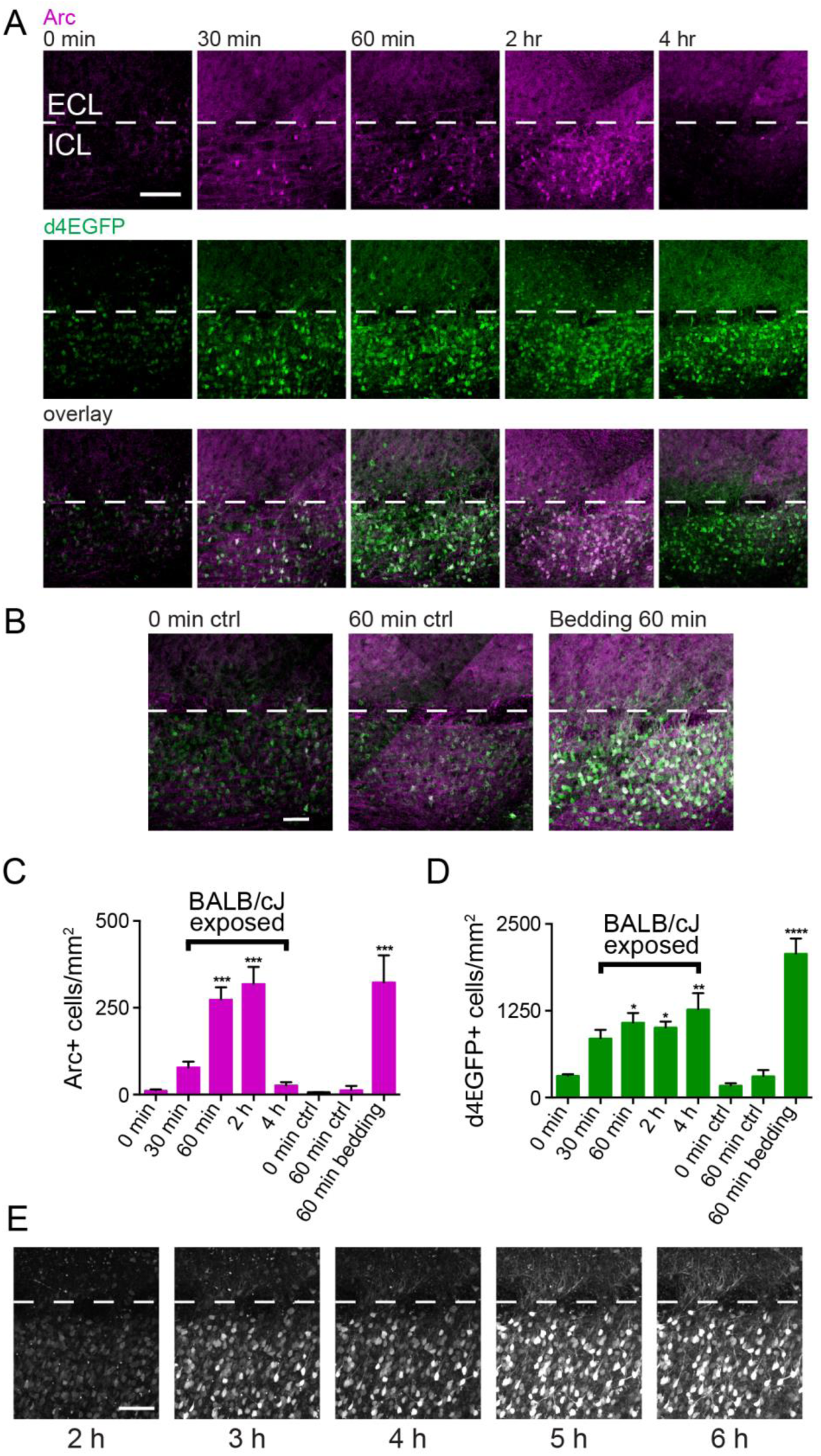
Arc protein expression peaks 2 hours after behavior and overlaps with d4EGFP expression in *Arc-d4EGFP*-BAC mice. ***A,*** Parasagittal AOB sections from resident mice perfused 0, 30, 60, 120, or 240 minutes after behavior, stained for both Arc protein and d4EGFP. The ICL is indicated by white dotted lines, posterior AOB is shown. Scale bar: 100 µm. ***B,*** Arc protein and d4EGFP expression in negative control mice that were perfused after 0 or 60 minutes in the behavioral setup, but were not exposed to a BALB/cJ intruder (left 2 panels). Arc protein and d4EGFP expression in a resident mouse that was exposed to soiled BALB/cJ bedding only in the behavioral setup, then perfused 60 minutes later. Scale bar: 50 µm. ***C,*** Quantification of Arc immunostaining across conditions. One-way ANOVA, F(7,11)=18.64, p<0.0001. ***D,*** Quantification of d4EGFP immunostaining across conditions. One-way ANOVA, F(7,11)=13.07, p=0.0002. For panels C and D, *, **, and ***, and **** represent p<0.05, 0.01, 0.001, and 0.0001, respectively, compared to the 0 min. controls (19 sections across 11 mice, ANOVA corrected for multiple comparisons using Dunnett’s method). ***E,*** Time course of d4EGFP expression in a single acute live slice visualized using 2-photon microscopy. Each image is a maximum z-projection of a 200 µm slice. Dissection occurred 1 hour after behavior. Scale bar: 100 µm.

**Figure 3:**
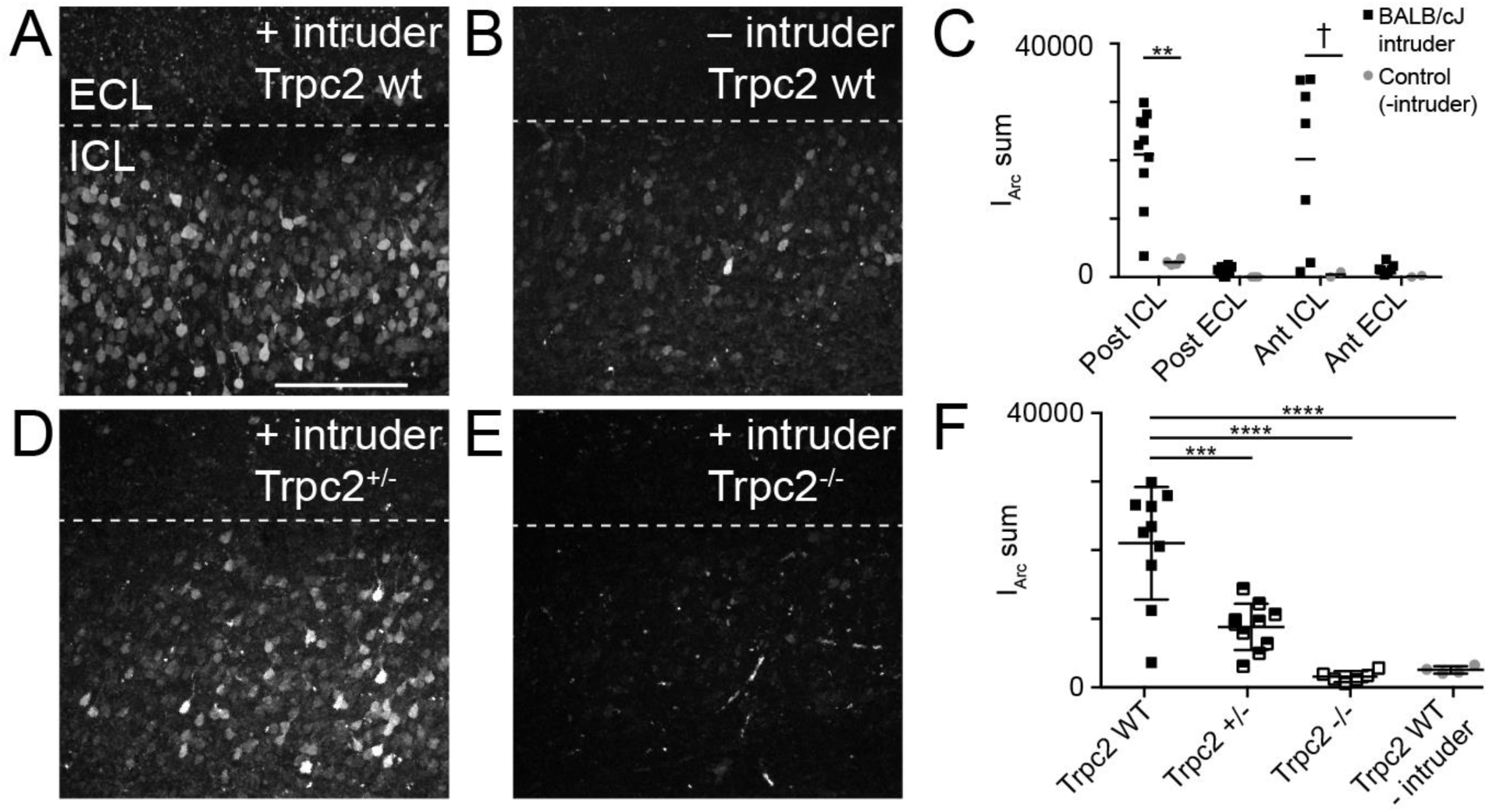
*Arc* expression is not upregulated after the resident-intruder assay in *Trpc2*^-/-^ mice. ***A,*** Maximum *z*-projection of an acute AOB slice taken from an *Arc-d4EGFP*-BAC, *Trpc2*^+/+^ resident. Posterior AOB is shown. Images were taken 4 hours after exposure to a BALB/cJ intruder. Scale bar: 200 µm. ***B,*** Representative z-projection taken from a control resident that was not exposed to an intruder. ***C,*** Quantification of summed fluorescence intensity across d4EGFP-expressing cells (I_Arc_ sum). Significance in each region determined using Wilcoxon-Mann-Whitney test. ** p<0.01; † p<0.1, Posterior ICL: n=9 mice, 10 slices (experimental), n=4 mice, 4 slices (control); Posterior ECL: n=9 mice, 10 slices (experimental), n=3 mice, 3 slices (control); Anterior ICL/ECL n=6 mice, 7 slices (experimental) n=2 mice, 2 slices (control). ***D,*** Representative z-projection taken from a *Trpc2*^+/-^ *Arc-d4EGFP*-BAC male resident. ***E,*** Representative z-projection taken from a *Trpc2*^-/-^ *Arc-d4EGFP*-BAC male resident. ***F,*** Quantification of I_Arc_ sum for the posterior ICL of all genotypes. *Trpc2*^+/-^ n=5 mice, 10 slices, *Trpc2*^-/-^ n=3 mice, 6 slices. One way ANOVA F(3,26)=22.75, p<0.0001. ***, and **** represent p<0.001, and 0.0001, respectively for indicated pairs (corrected for multiple comparisons using Tukey’s method).

**Figure 4:**
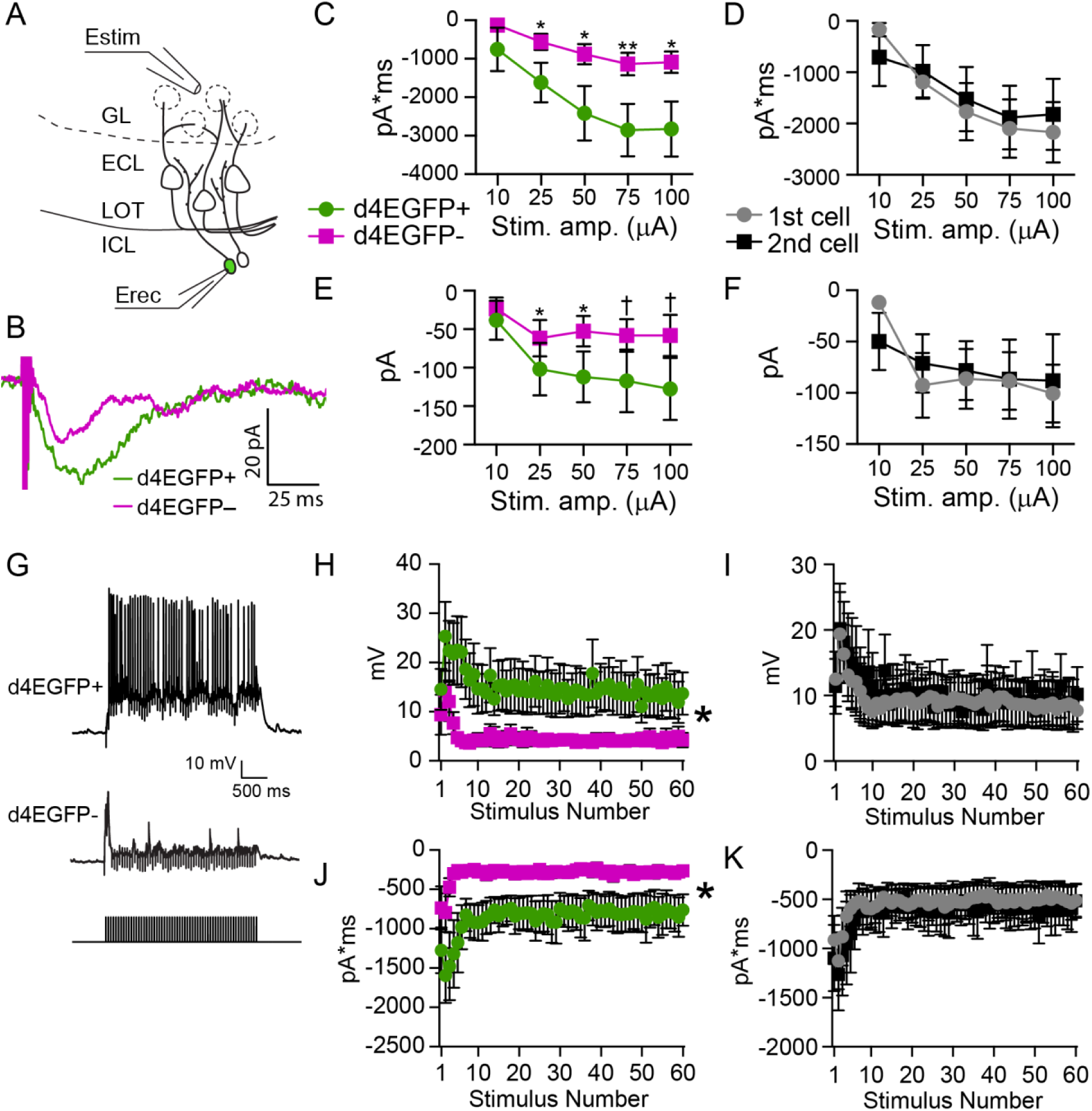
*Arc*-expressing IGCs show increased excitation by glomerular layer stimulation. ***A,*** Diagram illustrating experimental setup. Estim: theta glass stimulating electrode. Erec: recording electrode. ***B,*** Sample responses from a single, equal amplitude electrical stimulus from a d4EGFP+ cell and a nearby d4EGFP– cell from the same slice. ***C,*** Input-output curves showing charge transfer in response to a single stimulus pulse for d4EGFP+ and d4EGFP– cells. Paired, 2-tailed Student’s *t*-test. * p<0.05, ** p<0.01. ***D,*** Same data from panel C, grouped by the order in which the cells were recorded rather than d4EGFP expression. ***E,*** Input-output curves showing EPSC amplitude in response to a single pulse; data grouped by d4EGFP expression. Paired, 2-tailed Student’s *t*-test. † p<0.1 * p<0.05. ***F,*** Same data from panel E, grouped by the order in which cells were recorded. ***G,*** Sample traces from d4EGFP+ and d4EGFP– IGCs responding to 3 s, 20 Hz stimulation. ***H,*** Peak amplitude reached in response to each pulse of a 3 s, 20 Hz stimulus train. Repeated measures ANOVA, main effect of group (F(1,18)=4.51, p=0.048). ***I,*** Same data from panel H, grouped by the recording order. Repeated measures ANOVA, no main effect of group (F(1,18)=0.03, p=0.87) ***J,*** Total charge transfer in response to each pulse of the 3 s, 20 Hz stimulus train. Repeated measures ANOVA, main effect of group (F(1,18)=5.52, p=0.03). ***K,*** Same data from panel J, grouped by recording order. Repeated measures ANOVA, no main effect of group (F(1,18)=0.07, p=0.79). For all panels n=7 mice, 10 slices, 10 pairs.

**Figure 5:**
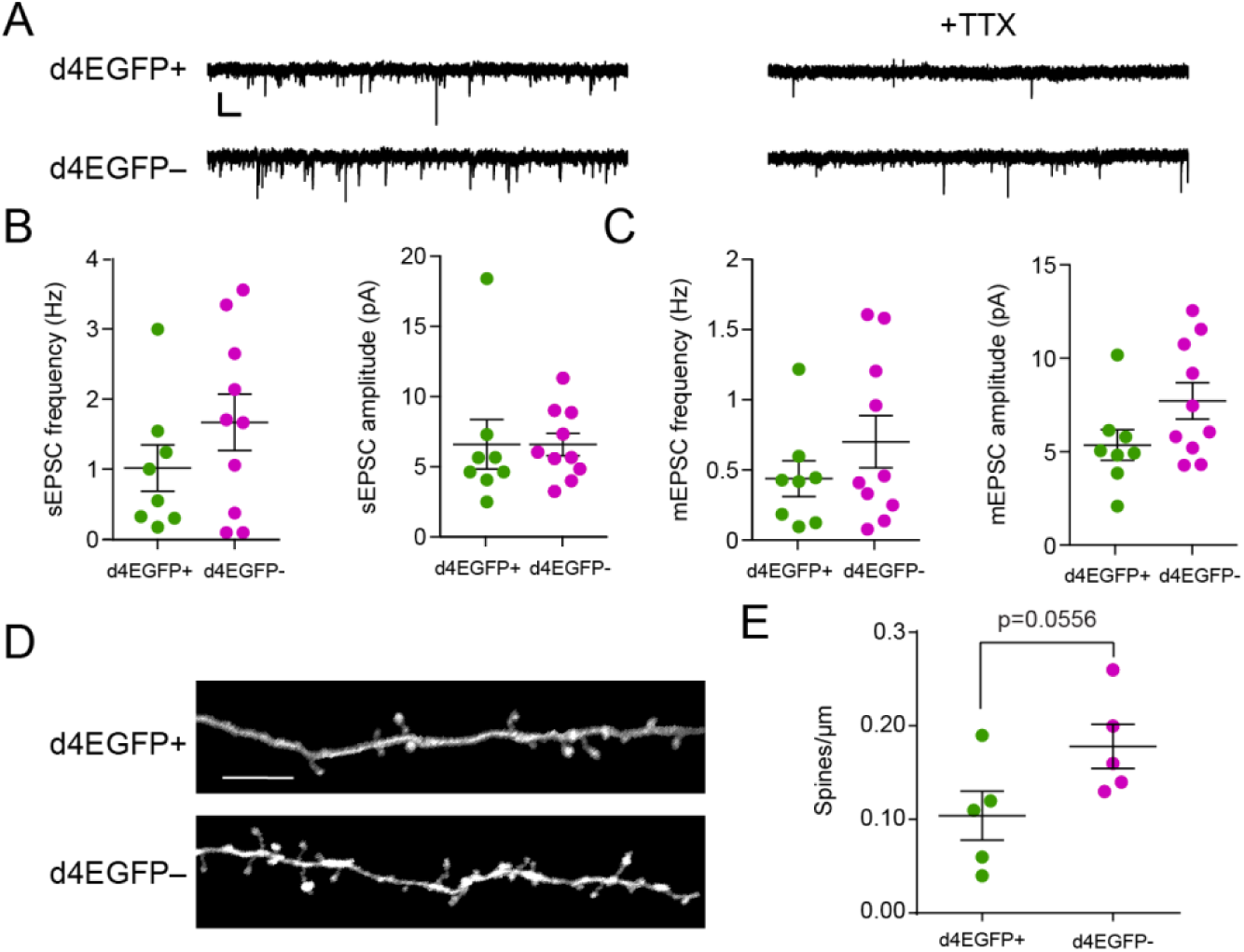
*Arc*-expressing IGCs do not display enhanced synaptic strength or number compared to non-expressing IGCs. ***A,*** Sample sEPSC and mEPSC traces from d4EGFP+ and d4EGFP– neurons. Scale bar: 10 pA, 500 ms. ***B,*** Spontaneous frequency and amplitude in d4EGFP+ IGCs. Significance determined using Wilcoxon-Mann-Whitney test (n.s., p>0.05). d4EGFP+ n=8, d4EGFP– n=10, n mice=10. ***C,*** Mini frequency and amplitude of d4EGFP+ and d4EGFP– IGCs. Significance determined using Wilcoxon-Mann-Whitney test (n.s., p>0.05). d4EGFP+ n=8, d4EGFP– n=10, n mice=10. ***D,*** Representative images showing dendritic spines on d4EGFP+ and d4EGFP– IGCs. Scale bar: 10 µm. ***E,*** Spines/um for d4EGFP+ and d4EGFP– neurons. Significance determined using Wilcoxon-Mann-Whitney test. n d4EGFP+ cells=5, n d4EGFP–cells=5, n mice=5.

### Electrophysiology

Whole cell patch-clamp recordings were made on d4EGFP+ and d4EGFP– IGCs during a window spanning 4-8 hours following behavior. Thin wall borosilicate glass electrodes with a tip resistance between 4 and 12 MΩ were filled with internal solution containing (in mM) 115 K-gluconate, 20 KCl, 10 HEPES, 2 EGTA, 2 MgATP, 0.3 Na_2_GTP, 10 Na phosphocreatine at pH 7.37. All recordings were amplified using a MultiClamp 700B amplifier (Molecular Devices) at 20 kHz and were digitized by a DigiData 1440 analog-digital converter controlled via pClamp 10.5 software (Molecular Devices, RRID:SCR_011323). Data were analyzed by Clampex 10.5 (Molecular Devices) and custom software written in MATLAB.

For glomerular stimulation experiments, a theta glass stimulating electrode was placed in the glomerular layer while recordings were made from IGCs. A series of 0.3 ms single pulses was used to construct input-output curves (Stimulus Isolator A365RC, World Precision Instruments), and the stimulation intensity for 20 Hz trains was the highest sub-saturating value from the input-output analysis. One d4EGFP+ and one d4EGFP– neuron were recorded per slice (one at a time). The d4EGFP+ neuron was recorded first in 6/10 pairs. Input-output curves were generated for each neuron, but the stimulation intensity used for both was chosen based on the first neuron recorded. Prior to MC recordings, we confirmed that stimulus conditions effectively recruited IGCs through direct IGC patch clamp recordings (23/24 recordings).

For mEPSC recordings, 5 minutes of baseline activity was recorded, then 1µM TTX was washed on for 5 minutes, and 5 minutes of post-drug activity was recorded.

Biocytin (3 mg/mL) and/or AlexaFluor568 (166 µM) was added to the internal solution to visualize dendritic arbors and spines. After filling, slices were fixed in 4% PFA in PBS for one hour, and then stored in PBS at 4°C.

To assess intrinsic electrophysiological features, we subjected patched AOB neurons to a series of current clamp and voltage clamp challenges. Immediately after achieving the whole cell configuration, each cell’s resting membrane potential (V_rest_) was measured in current clamp mode. To standardize measurements across cells with different V_rest_, we injected steady-state currents to maintain each cell’s membrane potential (V_m_) between -70 and -75 mV. Based on initial measurements of input resistance (R_input_), we empirically determined the amplitude of hyperpolarizing current that adjusted V_m_ by -50 mV (to ∼-125 mV). After determining this initial current injection amplitude, we generated a cell-specific 10-sweep Clampex protocol that applied increasingly depolarizing 0.5 s square current pulses, starting with the initial injection amplitude. For example, if the initial current injection was determined to be -100 pA, the 10-sweep protocol would have current injection increments of +20 pA (*i.e.*, -100 pA, -80 pA, -60 pA,…,+80 pA). If the initial depolarization was determined to be -125 pA, the protocol would include increments of +25 pA, etc. This strategy allowed us to objectively challenge cells with widely varying V_rest_ and R_input_. In voltage clamp, cells were initially held at -70 mV, and a series of 12 voltage command steps (0.5 s in duration) were applied that spanned -100 mV to +10 mV.

For each cell, both current clamp and voltage clamp protocols were applied up to 4 times, and all reported quantities represent the mean responses across repeated trials. Twenty-six specific intrinsic parameters were extracted from each cell using custom software written in MATLAB. A description of the parameters in Figure 6A and the formulas used to calculate them is presented in Table 1. I_H_ current ratio (Fig. 6G) was calculated using the formula

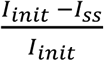

where *I*_*init*_ is the mean initial current (measured between 10 and 160 ms following the voltage pulse) and *I*_*ss*_ is the mean steady-state current (measured between 420 and 470 ms after the voltage pulse) generated by a voltage command step from -70 mV to -100 mV. EPSCs were automatically detected and later separated from noise using a custom computer assisted waveform-based event sorting program written in MATLAB (Hendrickson et al., 2008). EPSC decay was measured by calculating the best fit exponentially-decaying line for the decay period of the EPSC. Initial action potential rising slope was calculated by measuring the peak of the first derivative of voltage with respect to time (dV/dt). Threshold was defined as V_m_ at the time the dV/dt voltage reached 10% of its peak value (similar to Meeks et al., 2005). Membrane capacitance and input resistance were calculated according to current clamp-based multi-compartmental algorithms (Golowasch et al., 2009). Briefly, the voltage response of each cell to a hyperpolarizing current step was fit with a series of multi-exponential curves, and the best fit determined by identifying the solution with the lowest value of the Bayesian Information Criterion (to avoid over-fitting).

**Table 1.** Parameters used for multidimensional analysis.

| Shorthand | Description | Mode | Method | Reference |
| --- | --- | --- | --- | --- |
| Vrest | resting membrane potential | I Clamp | direct measurement | N/A |
| Cm | membrane capacitance | I Clamp | see reference | Golowasch, et al, 2009 |
| Rm | input resistance |  |  |  |
| Ncomps | number of model compartments | I Clamp | see reference |  |
| Ih_sag | hyperpolarization-induced depolarizing potential | I Clamp | $I_{H,sag} = V_{init} - V_{ss}$ | N/A |
| Ih_current | hyperpolarization-induced depolarizing current | V Clamp | $I_{H,curr} = I_{init} - I_{ss}$ | N/A |
| EPSC_freq | mean spontaneous EPSC frequency | V Clamp | direct measurement after waveform sorting | Hendrickson, et al, 2008 |
| EPSC_amp | mean spontaneous EPSC amplitude | V Clamp |  |  |
| EPSC_tau | mean spontaneous EPSC decay constant | V Clamp | least-squares exponential decay | N/A |
| S_freq_1 | Spiking frequency - Lv. 1 | I Clamp | direct measurement | N/A |
| S_freq_2 | Spiking frequency - Lv. 2 | I Clamp | direct measurement | N/A |
| S_freq_3 | Spiking frequency - Lv. 3 | I Clamp | direct measurement | N/A |
| S_freq_4 | Spiking frequency - Lv. 4 | I Clamp | direct measurement | N/A |
| S1_slope | Initial action potential rising slope | I Clamp | initial spike derivative peak | Meeks, et al, 2005 |
| S1_thresh | Initial action potential threshold | I Clamp | Vm @ 10% of initial spike derivative |  |
| S_accom_1 | Spike rate accommodation - Lv. 1 | I Clamp | $\frac{S_{init} - S_{final}}{S_{init} + S_{final}}$ | N/A |
| S_accom_2 | Spike rate accommodation - Lv. 2 | I Clamp |  |  |
| S_accom_3 | Spike rate accommodation - Lv. 3 | I Clamp |  |  |
| S_accom_4 | Spike rate accommodation - Lv. 4 | I Clamp |  |  |
| Na_curr_1 | Voltage-gated sodium current amplitude - Lv. 1 | V Clamp | direct measurement | N/A |
| Na_curr_2 | Voltage-gated sodium current amplitude - Lv. 2 | V Clamp | direct measurement | N/A |
| Na_curr_3 | Voltage-gated sodium current amplitude - Lv. 3 | V Clamp | direct measurement | N/A |
| Na_curr_4 | Voltage-gated sodium current amplitude - Lv. 4 | V Clamp | direct measurement | N/A |
| Na_curr_5 | Voltage-gated sodium current amplitude - Lv. 5 | V Clamp | direct measurement | N/A |
| K_curr_max | Voltage-gated potassium current - maximum | V Clamp | direct measurement | N/A |
| K_curr_diff | Non-inactivating voltage-gated potassium current | V Clamp | direct measurement | N/A |

**Figure 6:**
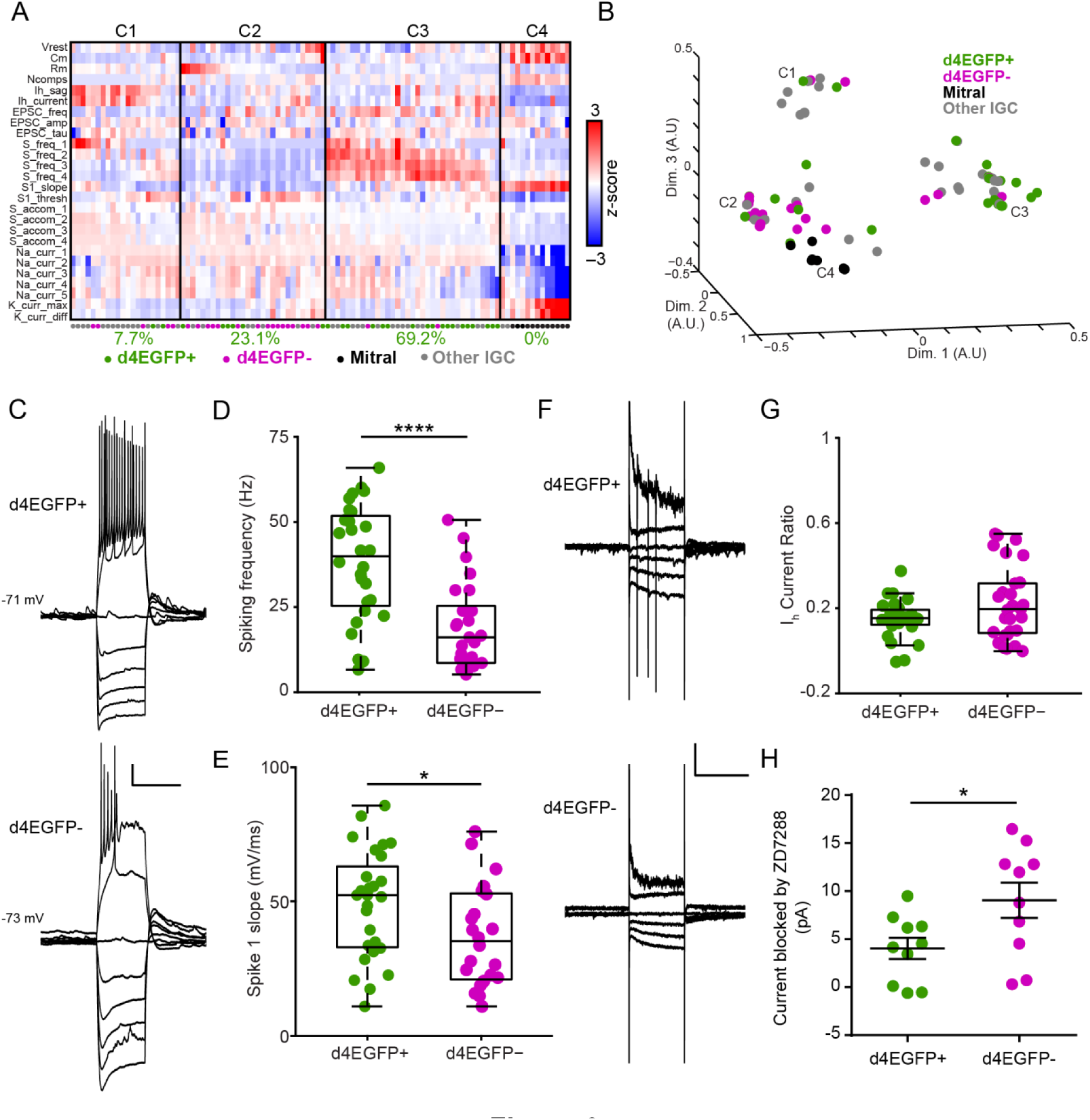
*Arc-* expressing IGCs are intrinsically more excitable than non-expressing IGCs. ***A,*** Colorized heat map representation of 26 intrinsic descriptors (rows) across 100 AOB neurons (columns) that were subjected to current-and voltage-clamp challenges. Solid vertical lines indicate divisions between identified clusters. Below each cluster is the percentage of all d4EGFP+ IGCs within that cluster. ***B,*** Multidimensional scaling of relative differences across all 26 dimensions from (A) into 3 dimensions. Each individual colored point indicates a cell, and C1-C4 refer to the cluster definitions in (A). For A and B, d4EGFP+ IGC n=26, d4EGFP– IGC n=23, n animals=15, other IGC n=39, mitral n=12, n animals=37. ***C,*** Sample traces for d4EGFP+ and d4EGFP– IGCs for current clamp ramp challenges. Scale bar: 10 mV, 500 ms. ***D,*** d4EGFP+ IGCs exhibit significantly increased spiking frequency when depolarized by a current injection (d4EGFP+ n=26, d4EGFP– n=23, Wilcoxon-Mann-Whitney test, p<0.0001). ***E,*** d4EGFP+ and d4EGFP– IGCs demonstrate increased maximal action potential slope (d4EGFP+ n=26, d4EGFP– n=23, Wilcoxon-Mann-Whitney test, p<0.05). ***F,*** Sample traces for d4EGFP+ and d4EGFP-IGCs for voltage clamp ramp challenges. Traces displayed show responses to being held at -100, -90, -80, -70, -40, and -20 mV. Scale bar: 50 pA, 500 ms. ***G,*** I_H_ current ratio for d4EGFP+ and d4EGFP– cells (d4EGFP+ n=26, d4EGFP– n=23, Wilcoxon-Mann-Whitney test, p=0.24). ***H,*** I_H_ subtracted currents following 10 µM ZD7288 application for d4EGFP+ and d4EGFP– IGCs (d4EGFP+ cells n,=10, d4EGFP–cells n=10, n mice=9, Wilcoxon-Mann-Whitney test, p<0.05).

## Results

### Arc expression is upregulated in posterior AOB IGCs following resident-intruder encounters

*Arc* is expressed in IGCs in rodents shortly after mating, an AOS-mediated behavior that induces pheromonal memory formation in females (Bruce and Parrott, 1960; Brennan et al., 1995; Matsuoka et al., 2002a; Matsuoka et al., 2003). If experience-dependent plasticity in the AOB is a more general response to chemosensory social events, we hypothesized that the social chemosensory encounters associated with male-male territorial aggression, which is AOS-mediated, would induce AOB plasticity. We used the resident-intruder paradigm, known to encourage AOS-dependent territorial aggression between males, as our behavioral challenge (Maruniak et al., 1986; Stowers et al., 2002). Adult **male** wild-type C57BL/6J **residents** were housed individually for one week and then exposed to adult male BALB/cJ intruders for 10 minutes. Arc protein expression was significantly increased 90 minutes after the behavior in AOB IGCs of resident males compared to controls (Wilcoxon-Mann-Whitney test, p<0.0001, Fig. 1A-D). Previous studies indicated that male-male resident-intruder encounters increase *Fos* expression in the posterior AOB (pAOB), which is selectively innervated by vomeronasal sensory neurons (VSNs) that express members of the V2R subfamily of vomeronasal receptors (Belluscio et al., 1999; Kumar et al., 1999; Rodriguez et al., 1999; Chamero et al., 2007). Consistent with these results, Arc protein expression after the resident-intruder paradigm was selectively upregulated in pAOB IGCs (Wilcoxon-Mann-Whitney test, p<0.001, Fig. 1E). These initial results show that *Arc* expression in IGCs following resident-intruder behavior is similar to *Arc* expression following mating behavior, and suggest that *Arc* expression occurs in an interneuron population that might increase MC inhibition after resident-intruder encounters.

### Arc-d4EGFP-BAC tools mark Arc-expressing IGCs in a time-and chemosignal-dependent manner

To identify *Arc*-expressing IGCs in live tissue, we utilized *Arc*-*d4EGFP*-BAC reporter mice (Grinevich et al., 2009). In these mice, a destabilized form of EGFP with a 4-hour half-life (d4EGFP) is expressed under control of the *Arc* promoter on a bacterial artificial chromosome, leaving endogenous *Arc* unperturbed. To assess the *Arc*-*d4EGFP*-BAC reporter, we used *Arc-d4EGFP*-BAC male mice as residents in the resident-intruder paradigm. We sacrificed animals at various time points following behavior (Fig. 2A) and compared Arc protein and d4EGFP immunostaining levels to controls (no intruder, BALB/cJ male soiled bedding only, Fig. 2A-D). Arc protein expression began rising by 30 minutes post-behavior, peaked 1-2 hours post-behavior and returned to baseline by 4 hours post-behavior (one-way ANOVA, F(7,11)=18.64, p<0.0001, n = 1-2 mice and 2-3 sections per condition, 11 mice and 19 sections overall, Fig. 2A, C). d4EGFP levels increased significantly by 1 hour post-behavior and remained elevated at 4 hours post-behavior (one-way ANOVA, F(7,11)=13.07, p=0.0002, Fig. 2 A, D). We observed strong colocalization between the Arc protein and d4EGFP signals, while some cells showed d4EGFP signal but no Arc protein in the cell soma. This effect could be explained by the fact that Arc protein is often localized to the dendrites (Shepherd and Bear, 2011), and that low levels of d4EGFP expression (indicating low level *Arc* transcription) were boosted by immunostaining.

Because d4EGFP immunostaining boosts very weak signals, we wanted to test whether d4EGFP expression alone was sufficient to identify *Arc*-expressing cells in living tissue. Live 2-photon imaging of acute AOB slices from *Arc*-*d4EGFP*-BAC residents revealed robust d4EGFP expression starting at 3 hours post-behavior and remaining strong until 6 hours post-behavior (Fig. 2E). Thus, unamplified d4EGFP signals lag behind the immunostaining time course, which is likely due to differences in antibody-amplified versus native d4EGFP signal. Importantly, the time window including the strongest behaviorally-driven d4EGFP expression was well-aligned with the time course of acute slice electrophysiological experiments, making the *Arc-d4EGFP*-BAC mice a strong tool for investigating the physiological properties of *Arc*-expressing IGCs.

Before we tested *Arc*-expressing IGC physiology, we wanted to determine whether d4EGFP expression required AOS sensory activation. This was important because different salient behaviors can activate centrifugal input into the entire olfactory bulb (Shipley et al., 1985; Brennan et al., 1990; Nunez-Parra et al., 2013; Rothermel et al., 2014; Oettl et al., 2016). Previous studies of AOB IEG expression indicated that chemosensory stimulation alone (soiled bedding) is sufficient to induce *Arc* expression and that social interaction increases this effect (Matsuoka et al., 2002b). Having confirmed that AOS chemosensory stimulation alone was sufficient to induce robust *Arc* expression (Fig. 2B-D), we wanted to next determine whether AOS activation was necessary for *Arc* expression. To accomplish this, we backcrossed *Arc-d4EGFP*-BAC reporter mice into a *Trpc2*^-/-^background. *Trpc2*, which is expressed in all vomeronasal sensory neurons and a small percentage of main olfactory sensory neurons, is required for proper chemosensory transduction (Omura and Mombaerts, 2014). Thus, *Trpc2*^-/-^ mice have severely disrupted vomeronasal chemosensory transduction and profound changes to AOS-mediated behaviors while retaining most main olfactory function (Stowers et al., 2002; Kimchi et al., 2007; Papes et al., 2010). We introduced *Arc-d4EGFP*-BAC, *Trpc2*^-/-^ resident males to BALB/cJ male intruders and imaged acutely-prepared live AOB slices using 2-photon microscopy 4 hours post-behavior, a time of robust d4EGFP expression after resident-intruder behaviors (Fig. 3). We quantified d4EGFP expression in the posterior ICL for *Trpc2*^+/+^, *Trpc2*^+/-^, and *Trpc2*^-/-^ animals exposed to intruders, as well as *Trpc2*^+/+^ residents that were left alone in an empty cage (no intruder) for 10 minutes (Fig. 3 A-F). Compared to the *Trpc2* wild-type, all other groups showed significantly reduced *d4E*GFP expression (one-way ANOVA F(3,26)=22.75, p<0.0001, Fig. 3F). This reduction occurred despite the fact that Trpc^-/-^ animals do not display deficits in olfactory social investigation (Stowers et al., 2002). Total d4EGFP intensity in *Trpc2*^-/-^ animals exposed to intruders matched that of *Trpc2*^+/+^ animals left alone in an empty cage, indicating that sensory input from VSNs is necessary for the elevated *Arc* expression in AOB IGCs (Fig. 3F). These results show that *Arc*-*d4EGFP*-BAC reporter mice label *Arc*-expressing IGCs in living AOB tissue, and show that VSN activation is both necessary and sufficient to induce AOB *Arc* expression after resident-intruder encounters.

### Arc-expressing IGCs respond strongly to glomerular stimulation

Previous work showed experience-dependent increases in inhibitory tone in the AOB following salient social behavior, suggesting that IGCs increase their inhibitory influence on MC output (Brennan et al., 1990; Brennan et al., 1995). If the IGCs expressing *Arc* after social chemosensory interactions contribute to AOB experience-dependent plasticity, we hypothesized that *Arc*-expressing IGCs would respond differently to AOB sensory input than non-expressing IGCs. To test this hypothesis, we made targeted whole cell patch clamp recordings from *Arc*-expressing (d4EGFP+) and non-expressing (d4EGFP–) IGCs in acute slices 4-8 hours after resident-intruder experiments (Fig. 4). To simulate sensory activity, we electrically stimulated VSN fibers in the AOB glomerular layer using theta-glass electrodes. Stimulating the glomerular layer induces glutamate release from VSN terminals and activates downstream MCs, which in turn activate IGCs at dendro-dendritic synapses (Fig. 4A). Thus, IGC activation resulting from VSN terminal stimulation is a di-synaptic effect. In voltage clamp, *Arc*-expressing IGCs showed increased single pulse EPSC charge transfer and peak current amplitude compared to non-expressing IGCs across stimulus intensities (paired, 2-tailed Student’s *t-* test, p<0.05, n=10 slices across 7 mice Fig. 4B-F). d4EGFP+ IGCs were recorded first in 6/10 experiments, so we investigated possible order effects by comparing these same responses based on the recording order (Fig. 4D, F). We found no differences in this comparison (paired, 2-tailed Student’s *t-* test, p>0.05), indicating that order effects cannot explain the increased excitability of d4EGFP+ IGCs. In order to evaluate the physiological relevance of these responses, we approximated strong MC activation by delivering a 3 s, 20 Hz train of stimuli to glomerular layer (matching MC firing rates measured during direct chemosensory stimulation of the VNO; Meeks et al., 2010). In current clamp, d4EGFP+ IGCS responded to trains with increased peak amplitudes, which were also sustained throughout the 3 second stimulus (repeated measures ANOVA, main effect of group F(1,18)=4.51, p=0.048, Fig. 4G-H). This effect was eliminated when the same data was organized based on recording order (repeated measures ANOVA, no main effect of group F(1,18)=0.03, p=0.87, Fig. 4I). In voltage clamp, the same d4EGFP+ IGCs responded to stimulus trains with significantly greater charge transfer than d4EGFP– IGCs and sustained this increase throughout the 3 seconds of stimulation (repeated measures ANOVA, main effect of group F(1,18)=5.52, p=0.03, Fig. 4J). As before, this effect was not due to an effect of recording order (repeated measures ANOVA, no main effect of group F(1,18)=0.07, p=0.79, Fig. 4K). These data indicate that *Arc*-expressing IGCs are more engaged by sensory input stimulation, which may reflect enhanced MC-IGC synaptic communication or increased IGC intrinsic excitability. These data also suggest that *Arc*-expressing IGCs may contribute to experience-dependent MC inhibition.

### Arc-expressing IGCs do not possess increased synapse strength or density

We hypothesized that the increased activation of *Arc*-expressing IGCs was due to a change in the number of MC-IGC synapses or a change in MC-IGC synaptic strength. To test these hypotheses, we recorded spontaneous and miniature EPSCs (sEPSCs and mEPSCs, respectively) from d4EGFP+ and d4EGFP– IGCs (Fig. 5). *Arc*-expressing IGCs did not exhibit a significant difference in sEPSC or mEPSC amplitude or frequency (Wilcoxon-Mann-Whitney test, p>0.05, Fig. 5A-C). In these experiments, biocytin was included in the recording pipette to allow *post hoc* analysis of dendrite morphology and dendritic spine density. Analysis of IGC dendrites showed a trend towards slightly lower spine densities in *Arc*-expressing IGCs compared to non-expressing IGCs (Wilcoxon-Mann-Whitney test, p=0.0556, Fig. 5D-E). Similarly there were no differences in average dendritic branch length (d4EGFP+: 485.3 ± 115.7 µm n=6, d4EGFP-: 365.5 ± 57.2 µm n=8, Wilcoxon-Mann-Whitney test, p=0.57), maximum dendritic branch length (d4EGFP+: 1813 ± 149.4 µm n=6, d4EGFP-: 1548 ± 231.4 µm n=8, Wilcoxon-Mann-Whitney test, p=0.41), or number of branches (d4EGFP+: 17.5 ± 4.75 branches n=6, d4EGFP-: 14.3 ± 1.89 branches n=8, Wilcoxon-Mann-Whitney test, p=0.59). Taken together, these results indicate that changes in IGC synaptic strength or number cannot explain the increased responses to glomerular stimulation in *Arc*-expressing IGCs.

### Arc-expressing IGCs possess enhanced intrinsic excitability

Lacking support for our initial hypothesis, we turned to an alternative, which was that *Arc*-expressing IGCs respond more strongly to VSN input stimulation due to changes in intrinsic properties. We utilized a systematic approach in order to assess possible intrinsic physiological differences in an unbiased way. We targeted whole-cell patch-clamp recordings to *Arc*-expressing and non-expressing IGCs 4-8 hours post-behavior and delivered a series of electrophysiological challenges in current and voltage clamp. Using automated algorithms (See Experimental Procedures), we quantified 26 physiological characteristics covering a variety of intrinsic properties (Table 1). To capture the differences between *Arc*-expressing and non-expressing IGCs, we performed cluster analysis on the 26-dimensional array of characteristics from 100 neurons that had undergone the same electrophysiological challenges (Fig. 6A-B). This collection of cells contained 26 d4EGFP+ and 23 d4EGFP– cells, along with 39 control IGCs from mice that had not undergone behavioral challenges and 12 MCs (used as a control population; Fig. 6A-B). 4 clusters were identified by this analysis. 69.2% of all d4EGFP+ IGCs were assigned to Cluster 3, where neurons exhibited sustained strong spiking activity in response to depolarization and lower levels of spike accommodation. Other intrinsic characteristics, including input resistance, resting membrane potential and action potential threshold, were highly variable across cells in this cluster, indicating that increased IGC firing frequency was not associated with a systematic change in these other properties.

In assessing individual intrinsic properties, few differed significantly between d4EGFP+ and d4EGFP– IGCs (Fig. 6C-H). The most noteworthy individual characteristic was an increase in spiking frequency in response to strong somatic depolarization in *Arc*-expressing IGCs (Wilcoxon-Mann-Whitney test, p<0.0001, Fig. 6D). d4EGFP+ IGCs demonstrated an increase in the maximal slope of initial action potentials (Wilcoxon-Mann-Whitney test, p<0.05, Fig. 6E), which was not a result of increased sodium current peak amplitudes (d4EGFP+: 2.33 ± 0.22 nA, d4EGFP–: 2.02 ± 0.14 nA, Wilcoxon-Mann-Whitney test, p=0.49) or spike threshold (d4EGFP+: -23.5 ± 1.5 mV, d4EGFP–: -23.0 ± 1.3 mV, Wilcoxon-Mann-Whitney test, p=0.53).

*Arc*-expressing IGCs also exhibited a slight decrease in hyperpolarization-activated cation (I_H_) currents (Fig. 6G), which we confirmed by measuring the currents blocked by the HCN channel antagonist ZD7288 (10 µM; Wilcoxon-Mann-Whitney test, p<0.05, Fig. 6H). Because IGCs have high input resistance (1.0 ± 0.17 GΩ, n = 54 IGCs), small amplitude I_H_ currents resulted in prominent depolarization (“sag” potential) measured at the soma (d4EGFP+: 5.81 ± 0.62 mV, d4EGFP-: 8.37 ± 0.90 mV, Wilcoxon-Mann-Whitney test, p=0.11). These basic I_H_ measurements were made after step hyperpolarization from a steady-state membrane potential near -70 mV, which may have obscured I_H_ conductances open near IGC resting potential. To measure the contributions of such conductances, we subtracted I_H_ sag potentials (reflecting conductances activated by the transition from ∼-70 to ∼-125 mV) from rebound depolarizations that occurred after the hyperpolarizing pulses (reflecting all I_H_ conductances). This analysis revealed a larger relative contribution of I_H_ conductances near resting potential in d4EGFP+ IGCs compared to d4EGFP– IGCs, suggesting a slight shift in the voltage-dependence of activation of HCN channels (d4EGFP+: 1.33 ± 0.57 mV, d4EGFP-: 0.23 ± 0.47 mV; Mann-Whitney-Wilcoxon test, p = 0.059). At face value, these I_H_ results were somewhat surprising. However, modulation of I_H_ has been reported in Arc-dependent plasticity elsewhere in the brain (Shah, 2014). In many cases downregulation of I_H_ has been associated with increased neuronal excitability (Poolos et al., 2002; Brager and Johnston, 2007; Campanac et al., 2008; Yi et al., 2016).

It may be possible that *Arc* is required for the expression of experience-dependent increases in IGC excitability. To investigate this hypothesis, we utilized *Arc*-d2EGFP knock-in/knock-out animals (Wang et al., 2006). In *Arc*^-/-^ animals, d2EGFP is expressed in neurons that would normally express *Arc.* We repeated our intrinsic electrophysiological assay in acute slices taken from Arc^-/-^ male residents. d2EGFP+ IGCs in *Arc*^-/-^ mice showed no differences in maximum spiking frequency (d2EGFP+:34.34 ± 4.64 Hz, n=16, d2EGFP-: 27.25 ± 4.38 Hz, n=12, p=0.30 Wilcoxon-Mann-Whitney test), I_H_ sag potential (d2EGFP+: 7.15 ± 0.71 mV, n=20, d2EGFP-: 7.80 ± 1.31 mV, n=14, Wilcoxon-Mann-Whitney test, p=0.45), or I_H_ currents (I_H_ current ratio d2EGFP+: 0.173 ± 0.025, n=17, d2EGFP–: 0.155 ± 0.029, n=14, Wilcoxon-Mann-Whitney test, p=0.80) compared to d2EGFP– IGCs from the same slices. These results suggest that *Arc* is required for the observed differences in excitability between *Arc-* expressing and non-expressing IGCs following male-male social interaction.

### MC responses to glomerular stimulation are suppressed following behavior

The increased excitability of *Arc*-expressing IGCs suggests that AOB MCs may experience stimulus-associated suppression following resident-intruder encounters. To test this hypothesis, we measured the responses of posterior AOB MCs to glomerular layer stimulation during the same 4-8 hour post-behavior time window used in IGC recordings (Fig. 7). MCs in current clamp were depolarized via steady-state current injections to -55 mV (just below action potential threshold), and exposed to 3 second, 20 Hz glomerular layer stimulation (Fig. 7B-C). We observed less evoked spiking activity in MCs in AOB slices taken from post-behavior resident males than no-intruder controls (repeated measures ANOVA, interaction between group and stimulus, F(59, 1298)=2.45, p<0.0001, n=13 cells, 4 mice for “intruder” group; n=9 cells, 4 mice for “no intruder” group, Fig. 7B-C). Voltage clamp experiments from these same cells held at -40 mV revealed suppressed net inward current during the stimulus trains (repeated measures ANOVA, main effect of group, F(1,22)=8.49, p=0.008, Fig. 7D). We observed no differences in net inward current when cells were held at -50 mV (nearer Cl^-^ _reversal_ potential), indicating that the results at -40 mV reflect the influence of increased outward currents (repeated measures ANOVA, no main effect of group F(1,22)=0.07, p=0.80). We did not observe increased MC intrinsic excitability post-behavior (MC max spiking frequency: 48.1 ± 2.5 Hz post-behavior, 40.2 ± 2.1 Hz control, Wilcoxon-Mann-Whitney test, p=0.13), but did involve a small decrease in I_H_ currents (normalized I_H_ current ratio 0.0342 ± 0.003 post-behavior, 0.047 ± 0.005, Wilcoxon-Mann-Whitney test, p=0.035). Importantly, these currents are much smaller than those observed in IGCs, consistent with recent observations in mitral cells (Gorin et al., 2016). These results confirm that male-male social chemosensory encounters are associated with subsequent MC suppression, consistent with an experience-dependent increase in IGC-MC inhibition.

**Figure 7:**
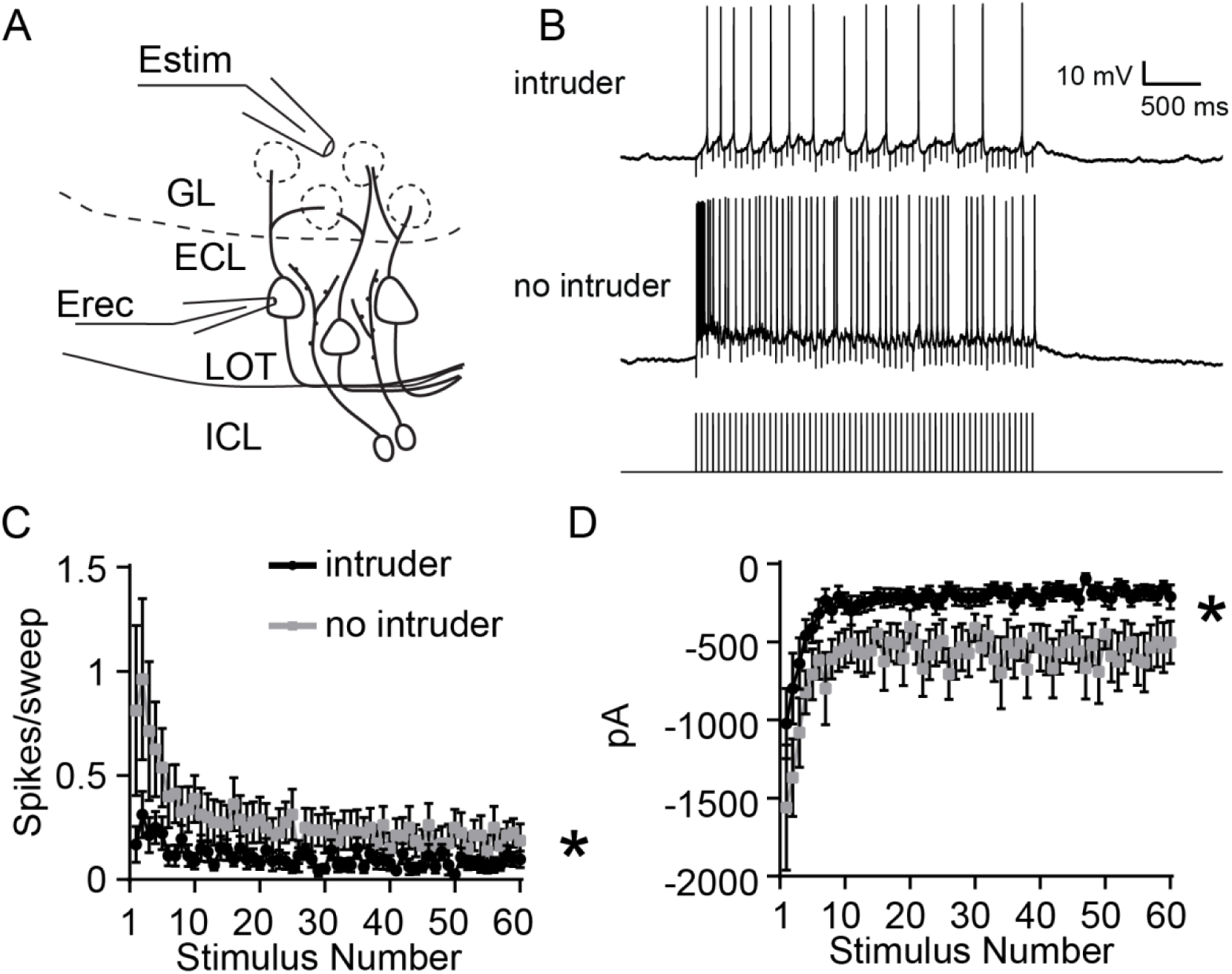
MCs show decreased excitation by glomerular stimulation following resident-intruder behavior. ***A,*** Diagram illustrating experimental setup. Estim: theta glass stimulating electrode. Erec: recording electrode. ***B,*** Sample MC responses to 3 s, 20 Hz GL stimulation from residents that interacted with an intruder and control residents (no intruder). ***C,*** Per-stimulus spike probability in response to each pulse of 3 s, 20 Hz stimulation while cell was artificially brought to the subthreshold potential of -55 mV. Repeated measures ANOVA, interaction between group and stimulus (F(59,1298)=2.45, p<.0001) Behavior: n cells=14, n mice=4. No behavior: n cells=10, n mice=4. ***D,*** Peak current amplitude in response to 3 s, 20 Hz stimulation while cell was held at -40 mV. Repeated measures ANOVA, main effect of group (F(1,22)=8.49, p=0.008). Behavior: n cells=13, n mice=4. No behavior: n cells=9, n mice=4.

## Discussion

The AOB remains a mysterious sensory circuit. It is often considered to be a relay station for information about environmental pheromones and kairomones, but there is ample evidence that the AOB also participates in experience-dependent plasticity (Brennan, 2009). Experience-dependent plasticity in the AOB has almost exclusively been studied in the female rodent AOB in the context of mating (Binns and Brennan, 2005; Brennan and Kendrick, 2006), but there are many other rodent behaviors that are strongly influenced by the AOS (Maruniak et al., 1986; Wysocki and Lepri, 1991; Stowers et al., 2002; Kimchi et al., 2007; Papes et al., 2010). The principal motivation for this work was a desire to learn more about experience-dependent AOB plasticity in the context of non-mating AOS-mediated behaviors.

We specifically chose to study the resident-intruder territorial aggression paradigm because it is a male-typical AOS-dependent behavior that induces robust IEG activation in the AOB (Maruniak et al., 1986; Wysocki and Lepri, 1991; Kumar et al., 1999; Stowers et al., 2002). *Arc* is upregulated in male and female rodent AOB IGCs following mating (Matsuoka et al., 2002a; Matsuoka et al., 2002b; Matsuoka et al., 2003), suggesting that *Arc*-expressing IGCs may underlie the increased AOB inhibition observed in this context (Kaba and Keverne, 1988; Brennan et al., 1990). Our results show that *Arc* is selectively transcribed and translated in posterior AOB IGCs of male mice following male-male social chemosensory encounters (Figs. 1-2). *Arc* upregulation in AOB IGCs after the resident-intruder assay requires AOS chemosensory signaling, indicating that this effect is not caused by brain processes (*e.g.*, centrifugal neuromodulation) associated with other sensory systems (Figs. 2-3).

The specific neurophysiological changes reported in Arc-expressing neurons in other brain regions vary widely and are often subtle (Wang et al., 2006; Ploski et al., 2008; Shepherd and Bear, 2011; Jakkamsetti et al., 2013). For example, *Arc*-deficient pyramidal neurons in visual cortex have an overall reduction in their orientation selectivity, but retain experience-dependent refinement of this selectivity (Wang et al., 2006). In the context of novel environment exploration, *Arc*-expressing hippocampal pyramidal neurons do not show outright synaptic depression, but instead are primed for mGluR-dependent LTD (Jakkamsetti et al., 2013). In the AOB, *Arc* upregulation is conspicuously absent in projecting MCs (Figs. 1-3). IGCs are physiologically and morphologically different than most of the principal cell types in which *Arc* has been studied. Specifically, IGCs are axonless and use reciprocal dendro-dendritic synapses to communicate with MCs (Jia et al., 1999; Taniguchi and Kaba, 2001). IGCs also experience significant cellular turnover in adulthood, and are replenished by adult-born neurons that migrate via the rostral migratory stream (Alvarez-Buylla and Garcia-Verdugo, 2002). The selective capacity of IGCs to upregulate *Arc* in response to social chemosensory experience suggests these interneurons may be primary drivers of AOB experience-dependent plasticity.

There are several genetic tools for labeling and manipulating *Arc*-expressing cells in living mouse brain tissue (Wang et al., 2006; Grinevich et al., 2009; Guenthner et al., 2013; Kawashima et al., 2014). We specifically chose *Arc-d4EGFP*-BAC reporter mice for our initial physiological experiments for two reasons. Firstly, endogenous *Arc* expression in these mice is unperturbed. Secondly, the half-life of the d4EGFP reporter allowed us to visualize *Arc*-expressing cells in acute slices for several hours following behavior. *Arc*-d2EGFP knock-in/knock-out mice remain a helpful tool for exploring the *Arc*-dependence of various forms of neuronal plasticity. However, experiments involving these mice use *Arc*-positive controls that are haploinsufficient for *Arc* and limit targeted physiology experiments to shorter time windows.

Previous studies of AOB IGC function used pharmacology to approximate the conditions present during salient social events (Araneda and Firestein, 2006; Smith et al., 2009; Taniguchi et al., 2012; Smith et al., 2015). These studies revealed important features of IGC neuromodulation, but did not investigate the cellular and synaptic changes that occur in IGCs activated by *bona fide* social behaviors. In *Arc-d4EGFP*-BAC mice, *Arc* expression in IGCs after resident-intruder encounters correlates with increased activation by sensory input (Fig. 4) and increased intrinsic excitability (Fig. 6). *Arc* expression has been associated with glutamate receptor trafficking in other contexts (Chowdhury et al., 2006; Shepherd et al., 2006; Waung et al., 2008). However, our data show that the increased network excitability of *Arc*-expressing IGCs is not related to an increase in EPSC frequency or amplitude, nor is there evidence of significant upregulation in the number of dendritic spines (Fig. 5). These results were somewhat surprising in light of previous work showing increased size of the postsynaptic density on IGC dendrites in female mice one day after mating, suggesting increased excitatory drive on IGCs (Matsuoka et al., 2004). However, our experiments took place both in a different behavioral context and at an earlier time point, which could indicate that different mechanisms or time courses are present following male-male social encounters. The increased excitability seen in *Arc*-expressing cells in d4EGFP mice was not observed in *Arc*^-/-^ mice from the *Arc*-d2EGFP strain. This indicates that *Arc* participates in the intrinsic differences in AOB IGCs seen after resident-intruder encounters. Collectively, these data suggest that *Arc* expression in IGCs contributes to an increased intrinsic capacity to respond to sensory input.

IGCs do not receive direct excitation from VSN terminals in the AOB glomerular layer, but are instead activated by glutamate release from MCs at reciprocal dendro-dendritic synapses (Jia et al., 1999; Taniguchi and Kaba, 2001). MCs do not demonstrate experience-dependent increases in intrinsic excitability following resident-intruder encounters, suggesting that the increased IGC activation is specific to the IGC postsynaptic response. However, it is also possible that a change in presynaptic function in the MCs providing input to *Arc*-expressing IGCs contributes to this effect. To approach this question will require tools that label both *Arc*-expressing IGCs and their connected MCs in living tissue.

In our investigation of the intrinsic differences between *Arc*-expressing and non-expressing cells, we employed methods aimed at objectively classifying cells based on the expression of 26 specific characteristics. *Arc*-expressing and non-expressing cells were segregated into clusters that differed in their capacity to sustain high frequency spiking (Fig. 6). The increase in excitability was not a result of changes to voltage gated sodium channel threshold, sodium current amplitudes, or macroscopic changes to voltage gated potassium currents. However, this analysis did reveal a trend for Arc-expressing IGCs to possess smaller I_H_ currents, which we confirmed pharmacologically (Fig. 6). HCN channel expression has been noted in the AOB, and in IGCs, but their specific role in AOB circuit function has not yet been determined (Hu et al., 2016). I_H_ is active at resting membrane potential and gives rise to rebound depolarization after relief from transient hyperpolarization (Robinson and Siegelbaum, 2003; Biel et al., 2009). At face value, the observed decrease in I_H_ is at odds with the increase in intrinsic excitability seen in *Arc*-expressing IGCs. However, several other studies have shown that plasticity-induced decreases in I_H_ are associated with increases in excitability (Poolos et al., 2002; Brager and Johnston, 2007; Campanac et al., 2008; Yi et al., 2016). One potential factor in the apparent excitatory influence of decreased I_H_ could be a shift in the voltage dependence of HCN channel activation, leading to an increased number of HCN channels open at rest. Overall, our results suggest that experience-dependent plasticity modulates IGC I_H_ currents, perhaps selectively in postsynaptic/dendritic structures. (Lorincz et al., 2002; Yi et al., 2016).

We confirmed that posterior MC activity is suppressed in AOB slices from resident mice following the resident-intruder assay (Fig. 7). This observation supports the hypothesis that experience-dependent upregulation of IGC excitability in *Arc*-expressing cells contributes to MC suppression. The effects were relatively modest, but were achieved despite lacking an experience-dependent marker to select for MCs that were active during the resident-intruder encounter. It is possible that this limitation precluded us from identifying other physiological changes in MCs that are induced following male-male social interaction.

The observed increases in IGC intrinsic excitability and MC suppression are present for at least 8 hours following behavior. This suggests that MC activation upon re-exposure to the same male during this time window would be decreased, which may result in a change in male-male social interactions. In the context of the Bruce Effect, pheromonal learning is AOB-dependent and can persist for many weeks (Brennan and Keverne, 1997). It may be the case that the AOS refines male-male social interactions over similar time courses, but the specific behavioral impacts and time courses of any effects remain to be elucidated. In sum, these data reveal that AOB experience-dependent plasticity involves *Arc* upregulation in IGCs, which results in increased MC inhibition through upregulation of IGC intrinsic excitability. Furthermore, our data show that inhibitory plasticity in the AOB occurs across social contexts and is a general feature of the AOS.

## Author Contributions

Conceptualization: H.L.C. and J.P.M.; Methodology: H.L.C., M.M., and J.P.M.; Software: J.P.M.; Formal Analysis: H.L.C. and J.P.M. Investigation: H.L.C., M.M., and J.P.M.; Resources: J.P.M, Data curation: H.L.C. and J.P.M; Writing: H.L.C. and J.P.M.; Visualization: H.L.C. and J.P.M., Supervision, Project Administration, and Funding Acquisition: J.P.M.

## Acknowledgements

We thank Jennifer Torres, Salma Ferdous, and Natasha Browder for technical support. We thank Kimberly Huber and Todd Roberts for critical feedback on the manuscript. This work was funded by the National Institute on Deafness and Other Communication Disorders and the National Institute on Drug Abuse of the National Institutes of Health under award numbers R00DC011780, R01DC015784 (JPM), and T32DA007290 (HLC). The content is solely the responsibility of the authors and does not necessarily represent the official views of the National Institutes of Health. Partial support was also provided by the National Science Foundation (Grant IOS-1451034; JPM, co-PI).

## REFERENCES

Alvarez-Buylla A, Garcia-Verdugo JM (2002) Neurogenesis in adult subventricular zone. J Neurosci 22:629–634.

Araneda RC, Firestein S (2006) Adrenergic enhancement of inhibitory transmission in the accessory olfactory bulb. J Neurosci 26:3292–3298.

Belluscio L, Koentges G, Axel R, Dulac C (1999) A map of pheromone receptor activation in the mammalian brain. Cell 97:209–220.

Biel M, Wahl-Schott C, Michalakis S, Zong X (2009) Hyperpolarization-activated cation channels: from genes to function. Physiological reviews 89:847–885.

Binns KE, Brennan PA (2005) Changes in electrophysiological activity in the accessory olfactory bulb and medial amygdala associated with mate recognition in mice. Eur J Neurosci 21:2529–2537.

Bolte S, Cordelieres FP (2006) A guided tour into subcellular colocalization analysis in light microscopy. Journal of microscopy 224:213–232.

Brager DH, Johnston D (2007) Plasticity of intrinsic excitability during long-term depression is mediated through mGluR-dependent changes in I(h) in hippocampal CA1 pyramidal neurons. J Neurosci 27:13926–13937.

Brennan P, Kaba H, Keverne EB (1990) Olfactory recognition: a simple memory system. Science 250:1223–1226.

Brennan PA (2009) Outstanding issues surrounding vomeronasal mechanisms of pregnancy block and individual recognition in mice. Behav Brain Res 200:287–294.

Brennan PA, Keverne EB (1997) Neural mechanisms of mammalian olfactory learning. Prog Neurobiol 51:457–481.

Brennan PA, Kendrick KM (2006) Mammalian social odours: attraction and individual recognition. Philos Trans R Soc Lond B Biol Sci 361:2061–2078.

Brennan PA, Kendrick KM, Keverne EB (1995) Neurotransmitter release in the accessory olfactory bulb during and after the formation of an olfactory memory in mice. Neuroscience 69:1075–1086.

Bruce HM (1959) An exteroceptive block to pregnancy in the mouse. Nature 184:105.

Bruce HM, Parrott DM (1960) Role of olfactory sense in pregnancy block by strange males. Science 131:1526.

Campanac E, Daoudal G, Ankri N, Debanne D (2008) Downregulation of dendritic I(h) in CA1 pyramidal neurons after LTP. J Neurosci 28:8635–8643.

Chamero P, Marton TF, Logan DW, Flanagan K, Cruz JR, Saghatelian A, Cravatt BF, Stowers L (2007) Identification of protein pheromones that promote aggressive behaviour. Nature 450:899–902.

Chowdhury S, Shepherd JD, Okuno H, Lyford G, Petralia RS, Plath N, Kuhl D, Huganir RL, Worley PF (2006) Arc/Arg3.1 interacts with the endocytic machinery to regulate AMPA receptor trafficking. Neuron 52:445–459.

Ferrero DM, Moeller LM, Osakada T, Horio N, Li Q, Roy DS, Cichy A, Spehr M, Touhara K, Liberles SD (2013) A juvenile mouse pheromone inhibits sexual behaviour through the vomeronasal system. Nature 502:368–371.

Golowasch J, Thomas G, Taylor AL, Patel A, Pineda A, Khalil C, Nadim F (2009) Membrane capacitance measurements revisited: dependence of capacitance value on measurement method in nonisopotential neurons. J Neurophysiol 102:2161–2175.

Gorin M, Tsitoura C, Kahan A, Watznauer K, Drose DR, Arts M, Mathar R, O’Connor S, Hanganu-Opatz IL, Ben-Shaul Y, Spehr M (2016) Interdependent Conductances Drive Infraslow Intrinsic Rhythmogenesis in a Subset of Accessory Olfactory Bulb Projection Neurons. J Neurosci 36:3127–3144.

Grinevich V, Kolleker A, Eliava M, Takada N, Takuma H, Fukazawa Y, Shigemoto R, Kuhl D, Waters J, Seeburg PH, Osten P (2009) Fluorescent Arc/Arg3.1 indicator mice: a versatile tool to study brain activity changes in vitro and in vivo. J Neurosci Meth 184:25–36.

Guenthner CJ, Miyamichi K, Yang HH, Heller HC, Luo L (2013) Permanent genetic access to transiently active neurons via TRAP: targeted recombination in active populations. Neuron 78:773–784.

Guthrie K, Rayhanabad J, Kuhl D, Gall C (2000) Odors regulate Arc expression in neuronal ensembles engaged in odor processing. Neuroreport 11:1809–1813.

Hendrickson RC, Krauthamer S, Essenberg JM, Holy TE (2008) Inhibition shapes sex selectivity in the mouse accessory olfactory bulb. J Neurosci 28:12523–12534.

Hu R, Ferguson KA, Whiteus CB, Meijer DH, Araneda RC (2016) Hyperpolarization-Activated Currents and Subthreshold Resonance in Granule Cells of the Olfactory Bulb. eNeuro 3.

Jakkamsetti V, Tsai NP, Gross C, Molinaro G, Collins KA, Nicoletti F, Wang KH, Osten P, Bassell GJ, Gibson JR, Huber KM (2013) Experience-induced Arc/Arg3.1 primes CA1 pyramidal neurons for metabotropic glutamate receptor-dependent long-term synaptic depression. Neuron 80:72–79.

Jia C, Chen WR, Shepherd GM (1999) Synaptic organization and neurotransmitters in the rat accessory olfactory bulb. J Neurophysiol 81:345–355.

Kaba H, Keverne EB (1988) The effect of microinfusions of drugs into the accessory olfactory bulb on the olfactory block to pregnancy. Neuroscience 25:1007–1011.

Kawashima T, Okuno H, Bito H (2014) A new era for functional labeling of neurons: activity-dependent promoters have come of age. Frontiers in neural circuits 8:37.

Kawashima T, Kitamura K, Suzuki K, Nonaka M, Kamijo S, Takemoto-Kimura S, Kano M, Okuno H, Ohki K, Bito H (2013) Functional labeling of neurons and their projections using the synthetic activity-dependent promoter E-SARE. Nat Methods 10:889–895.

Kim Y, Venkataraju KU, Pradhan K, Mende C, Taranda J, Turaga SC, Arganda-Carreras I, Ng L, Hawrylycz MJ, Rockland KS, Seung HS, Osten P (2015) Mapping social behavior-induced brain activation at cellular resolution in the mouse. Cell reports 10:292–305.

Kimchi T, Xu J, Dulac C (2007) A functional circuit underlying male sexual behaviour in the female mouse brain. Nature 448:1009–1014.

Kumar A, Dudley CA, Moss RL (1999) Functional dichotomy within the vomeronasal system: distinct zones of neuronal activity in the accessory olfactory bulb correlate with sex-specific behaviors. The Journal of neuroscience: the official journal of the Society for Neuroscience 19:RC32.

Larriva-Sahd J (2008) The accessory olfactory bulb in the adult rat: a cytological study of its cell types, neuropil, neuronal modules, and interactions with the main olfactory system. J Comp Neurol 510:309–350.

Leinders-Zufall T, Ishii T, Chamero P, Hendrix P, Oboti L, Schmid A, Kircher S, Pyrski M, Akiyoshi S, Khan M, Vaes E, Zufall F, Mombaerts P (2014) A family of nonclassical class I MHC genes contributes to ultrasensitive chemodetection by mouse vomeronasal sensory neurons. J Neurosci 34:5121–5133.

Liberles SD (2014) Mammalian pheromones. In, pp 151–175.

Longair MH, Baker DA, Armstrong JD (2011) Simple Neurite Tracer: open source software for reconstruction, visualization and analysis of neuronal processes. Bioinformatics 27:2453–2454.

Lorincz A, Notomi T, Tamas G, Shigemoto R, Nusser Z (2002) Polarized and compartment-dependent distribution of HCN1 in pyramidal cell dendrites. Nat Neurosci 5:1185–1193.

Madisen L, Zwingman TA, Sunkin SM, Oh SW, Zariwala HA, Gu H, Ng LL, Palmiter RD, Hawrylycz MJ, Jones AR, Lein ES, Zeng H (2010) A robust and high-throughput Cre reporting and characterization system for the whole mouse brain. Nature neuroscience 13:133–140.

Maruniak JA, Wysocki CJ, Taylor JA (1986) Mediation of male mouse urine marking and aggression by the vomeronasal organ. Physiol Behav 37:655–657.

Matsuoka M, Yamagata K, Sugiura H, Yoshida-Matsuoka J, Norita M, Ichikawa M (2002a) Expression and regulation of the immediate-early gene product Arc in the accessory olfactory bulb after mating in male rat. Neuroscience 111:251–258.

Matsuoka M, Yoshida-Matsuoka J, Sugiura H, Yamagata K, Ichikawa M, Norita M (2002b) Mating behavior induces differential Arc expression in the main and accessory olfactory bulbs of adult rats. Neurosci Lett 335:111–114.

Matsuoka M, Yoshida-Matsuoka J, Yamagata K, Sugiura H, Ichikawa M, Norita M (2003) Rapid induction of Arc is observed in the granule cell dendrites in the accessory olfactory bulb after mating. Brain research 975:189–195.

Matsuoka M, Kaba H, Moriya K, Yoshida-Matsuoka J, Costanzo RM, Norita M, Ichikawa M (2004) Remodeling of reciprocal synapses associated with persistence of long-term memory. Eur J Neurosci 19:1668–1672.

Meeks JP, Jiang X, Mennerick S (2005) Action potential fidelity during normal and epileptiform activity in paired soma-axon recordings from rat hippocampus. J Physiol 566:425–441.

Meeks JP, Arnson HA, Holy TE (2010) Representation and transformation of sensory information in the mouse accessory olfactory system. Nat Neurosci 13:723–730.

Nunez-Parra A, Maurer RK, Krahe K, Smith RS, Araneda RC (2013) Disruption of centrifugal inhibition to olfactory bulb granule cells impairs olfactory discrimination. Proc Natl Acad Sci U S A 110:14777–14782.

Oettl LL, Ravi N, Schneider M, Scheller MF, Schneider P, Mitre M, da Silva Gouveia M, Froemke RC, Chao MV, Young WS, Meyer-Lindenberg A, Grinevich V, Shusterman R, Kelsch W (2016) Oxytocin Enhances Social Recognition by Modulating Cortical Control of Early Olfactory Processing. Neuron 90:609–621.

Omura M, Mombaerts P (2014) Trpc2-expressing sensory neurons in the main olfactory epithelium of the mouse. Cell reports 8:583–595.

Pankevich DE, Baum MJ, Cherry JA (2004) Olfactory sex discrimination persists, whereas the preference for urinary odorants from estrous females disappears in male mice after vomeronasal organ removal. The Journal of neuroscience: the official journal of the Society for Neuroscience 24:9451–9457.

Papes F, Logan DW, Stowers L (2010) The vomeronasal organ mediates interspecies defensive behaviors through detection of protein pheromone homologs. In: Cell, pp 692–703.

Perez-Gomez A, Bleymehl K, Stein B, Pyrski M, Birnbaumer L, Munger SD, Leinders-Zufall T, Zufall F, Chamero P (2015) Innate Predator Odor Aversion Driven by Parallel Olfactory Subsystems that Converge in the Ventromedial Hypothalamus. Curr Biol 25:1340–1346.

Plath N et al. (2006) Arc/Arg3.1 is essential for the consolidation of synaptic plasticity and memories. Neuron 52:437–444.

Ploski JE, Pierre VJ, Smucny J, Park K, Monsey MS, Overeem KA, Schafe GE (2008) The activity-regulated cytoskeletal-associated protein (Arc/Arg3.1) is required for memory consolidation of pavlovian fear conditioning in the lateral amygdala. J Neurosci 28:12383–12395.

Poolos NP, Migliore M, Johnston D (2002) Pharmacological upregulation of h-channels reduces the excitability of pyramidal neuron dendrites. Nat Neurosci 5:767–774.

Robinson RB, Siegelbaum SA (2003) Hyperpolarization-activated cation currents: from molecules to physiological function. Annual review of physiology 65:453–480.

Rodriguez I, Feinstein P, Mombaerts P (1999) Variable patterns of axonal projections of sensory neurons in the mouse vomeronasal system. Cell 97:199–208.

Rothermel M, Carey RM, Puche A, Shipley MT, Wachowiak M (2014) Cholinergic inputs from Basal forebrain add an excitatory bias to odor coding in the olfactory bulb. J Neurosci 34:4654–4664.

Scalia F, Winans SS (1975) The differential projections of the olfactory bulb and accessory olfactory bulb in mammals. J Comp Neurol 161:31–55.

Shah MM (2014) Cortical HCN channels: function, trafficking and plasticity. J Physiol 592:2711–2719.

Shakhawat AM, Gheidi A, Hou Q, Dhillon SK, Marrone DF, Harley CW, Yuan Q (2014) Visualizing the engram: learning stabilizes odor representations in the olfactory network. J Neurosci 34:15394–15401.

Shepherd JD, Bear MF (2011) New views of Arc, a master regulator of synaptic plasticity. Nat Neurosci 14:279–284.

Shepherd JD, Rumbaugh G, Wu J, Chowdhury S, Plath N, Kuhl D, Huganir RL, Worley PF (2006) Arc/Arg3.1 mediates homeostatic synaptic scaling of AMPA receptors. Neuron 52:475–484.

Shipley MT, Halloran FJ, de la Torre J (1985) Surprisingly rich projection from locus coeruleus to the olfactory bulb in the rat. Brain Res 329:294–299.

Smith RS, Weitz CJ, Araneda RC (2009) Excitatory actions of noradrenaline and metabotropic glutamate receptor activation in granule cells of the accessory olfactory bulb. J Neurophysiol 102:1103–1114.

Smith RS, Hu R, DeSouza A, Eberly CL, Krahe K, Chan W, Araneda RC (2015) Differential Muscarinic Modulation in the Olfactory Bulb. J Neurosci 35:10773–10785.

Stowers L, Holy TE, Meister M, Dulac C, Koentges G (2002) Loss of sex discrimination and male-male aggression in mice deficient for TRP2. Science 295:1493–1500.

Taniguchi M, Kaba H (2001) Properties of reciprocal synapses in the mouse accessory olfactory bulb. Neuroscience 108:365–370.

Taniguchi M, Yokoi M, Shinohara Y, Okutani F, Murata Y, Nakanishi S, Kaba H (2012) Regulation of synaptic currents by mGluR2 at reciprocal synapses in the mouse accessory olfactory bulb. Eur J Neurosci.

Vazdarjanova A, Ramirez-Amaya V, Insel N, Plummer TK, Rosi S, Chowdhury S, Mikhael D, Worley PF, Guzowski JF, Barnes CA (2006) Spatial exploration induces ARC, a plasticity-related immediate-early gene, only in calcium/calmodulin-dependent protein kinase II-positive principal excitatory and inhibitory neurons of the rat forebrain. J Comp Neurol 498:317–329.

Vousden DA, Epp J, Okuno H, Nieman BJ, van Eede M, Dazai J, Ragan T, Bito H, Frankland PW, Lerch JP, Henkelman RM (2015) Whole-brain mapping of behaviourally induced neural activation in mice. Brain structure & function 220:2043–2057.

Wang KH, Majewska A, Schummers J, Farley B, Hu C, Sur M, Tonegawa S (2006) In vivo two-photon imaging reveals a role of arc in enhancing orientation specificity in visual cortex. Cell 126:389–402.

Waung MW, Pfeiffer BE, Nosyreva ED, Ronesi JA, Huber KM (2008) Rapid translation of Arc/Arg3.1 selectively mediates mGluR-dependent LTD through persistent increases in AMPAR endocytosis rate. Neuron 59:84–97.

Wysocki CJ, Lepri JJ (1991) Consequences of removing the vomeronasal organ. J Steroid Biochem Mol Biol 39:661–669.

Yi F, Danko T, Botelho SC, Patzke C, Pak C, Wernig M, Sudhof TC (2016) Autism-associated SHANK3 haploinsufficiency causes Ih channelopathy in human neurons. Science 352:aaf2669.

